# Gene-centric intra- and inter-clade recombination in a context of *Esche-richia coli* subpopulations

**DOI:** 10.1101/122713

**Authors:** Yu Kang, Xing Shi, Lina Yuan, Yanan Chu, Fei Chen, Zilong He, Zhancheng Gao, Xinmiao Jia, Qiang Lin, Qin Ma, Jian Wang, Rongrong Fu, Jiayan Wu, Jingfa Xiao, Songnian Hu, Jun Yu

**Affiliations:** CAS Key Laboratory of Genome Sciences and Information, Beijing Institute of Genomics, Chinese Academy of Sciences, 100101, Beijing, PR China.; University of Chinese Academy of Sciences, Beijing 100190, PR China.; Department of Respiratory & Critical Care Medicine, Peking University People’s Hospital, Beijing, 100044, PR China; Central Research Laboratory, Peking Union Medical College Hospital, Peking Union Medical College & Chinese Academy of Medical Sciences, Beijing 100730, P.R.China; Department of Agronomy, Horticulture, and Plant Science, South Dakota State University, Brookings, SD, 57007, USA.

## Abstract

Recombination is one of the most important mechanisms of prokaryotic species evolution but its exact roles are still in debate. Here we try to infer genome-wide recombination events within a species uti-lizing a dataset of 104 complete genomes of *Escherichia coli* from diverse origins, among which 45 from world-wide animal-hosts are in-house sequenced using SMRT (single-molecular real time) technology.Two major clades are identified based on evidences of ecological and physiological characteristics, as well as distinct genomic features implying scarce inter-clade genetic exchange. By comparing the synteny of identical fragments genome-widely searched for each genome pair, we achieve a fine-scale map of re-combination within the population. The recombination is rather extensive within clade, which is able to break linkages between genes but does not interrupt core genome framework and primary metabolic port-folios possibly due to natural selection for physiological compatibility and ecological fitness. Meanwhile,the recombination between clades declines drastically as the phylogenetic distance increases, generally 10-fold reduced than those of the intra-clade, which establishes genetic barrier between clades. These empirical data of recombination suggest its critical role in the early stage of speciation, where recombina-tion rate differs according to phylogentic distance. The extensive intra-clade recombination coheres sister strains into a quasi-sexual group and optimizes genes or alleles to streamline physiological activities,whereas shapely declined inter-clade recombination split the population into clades adaptive to divergent ecological niches.

**Significance Statement:** Roles of recombination in species evolution have been debated for decades due to difficulties in inferring recombination events during the early stage of speciation, especially when recombination is always complicated by frequent gene transfer events of bacterial genomes. Based on 104 high-quality complete *E. coli* genomes, we infer gene-centric dynamics of recombination in the formation of two *E. coli* clades or subpopulations, and recombination is found to be rather intensive in a within-clade fashion, which forces them to be quasi-sexual. The recombination events can be mapped among individual genomes in the context of genes and their variations; decreased between-clade and increased intra-clade recombination engender a genetic barrier that further encourages clade-specific secondary metabolic portfolios for better environmental adaptation. Recombination is thus a major force that accelerates bacterial evolution to fit ecological diversity.

## Main Text

### Introduction

Roles of recombination in bacterial evolution have been debated for decades (1, 2). Asexual bacteria are believed to rarely recombine their genomes when passing genetic information to progenies, and a cluster of genes is capable of taking over the entire population if they happen to be linked with an advantageous gene through genetic hitchhiking, leading to an effect of genome-wide selective sweep as proposed in an “ecotype model”(3, 4). The theory is supported by a recent time-series metagenomics research (5), whereas other studies reveal the counterpart where recombination breaks the linkage and leaves only the adaptive gene to sweep a population (6, 7). The issue comes that to what extent recombination actually breaks such genetic linkages and undermines the theory of genome sweep. Another issue relevant to recombination is its contradictory roles in speciation, where recombination produces inconsistent signals in clonal phylogenies (8) in one hand and hijacks similar strains into a separating subpopulation (1, 9) in the other. Recent studies have postulated extensive recombination in bacterial populations even to a quasi-sexual status(6). Will extensive recombination interrupt primary metabolic portfolios or genome organization frameworks (GOF) (10), and lead to puzzled or discordant phylogenies? These issues can be readily settled by resolving recombination events in a genome-scale and at single gene level within a bacterial population where subpopulations are recently diverged, making a fine-scale and panoramic display of the recombination dynamics in early speciation. However, it is still unachievable due to the difficulties in inferring recombination events between closely related strains (11).

Here, we investigate the species *E. coli*, an excellent model for studying genotype-phenotype relationships and molecular evolution mechanisms due to its collective knowledge, ecological diversity, and availability of high-quality genomes and their annotations. Pangenomic studies have revealed great genome variability and intensive recombination in *E. coli,* which underlie discordance between phylogenies of various orthologous genes (12). To generate recombination dynamics genome-wide and in a fine-scale, we start with 45 in-house genome assemblies from animal hosts, using SMRT (Single-molecule real-time sequencing) technology, and also put together a collection of 104 complete genome sequences of strains from diverse geographic and host range from public databases. Based on genome-based phylogenies coupled with experimentation and detailed genetic analysis, we further classify *E. coli* isolates into two major clades or subpopulations, *Vigorous* for its fast movement and *Sluggish* for its slow movement, abbreviated as Vig and Slu. Taking advantage of our complete genome assemblies, as well as their precise synteny of identical fragments and genes, we are able to infer recombination events for each genome pair and to calculate the mobility (transfer among isolates within and across clades) of each gene, which are used to address: (1) In what degree and intensity does recombination happen within a species and its subpopulations? (2) What are molecular and physiological bases of the early separation of clades? (3) Does recombination disrupt GOF and metabolic portfolios of a host strain?

### Results

Our study begins with a thorough collection of 202 *E. coli* strains of animal hosts with broad diversity in geography, climate, and host range by field scientists, and the strains have been served as representation of the species for detailed taxonomic characterization (13). Of this collection, 45 isolates which represent maximal genetic heterogeneity and biodiversity in terms of host diet, geographic distribution (Fig. S1) and MLST clustering (Fig. S2), are selected for genome sequencing with both Illunima and Pac-Bio RSII platform; and all genomes are finished to a single circular chromosome.

For sequence analysis, we retrieve 59 complete genomes of human-host strains together with 7 draft genomes of environmental strains (14) from the public databases. Based on the 111 genomes, we extract the core genome of *E. coli,* which has collectively 1,095 genes in a total length of 1.05 Mb. The maximal-likelihood phylogeny based on the core genome indicates clear separation between host-related and environmental strains, whereas human- and animal-host strains appear entangled (Fig.1). Most host-related strains cluster into two major clades; the larger one contains 56 strains of phylotype A and B1, whereas the smaller one is composed of 31 phylotype B2 strains. The rest 17 strains, mainly in phylotype D and E, form several minor clades and are more closely positioned to the environment strains. Overall, this clustering pattern is largely in congruence with their phylotypes and previous reports (15, 16). As for the two major clades, although they both contain commensals and pathogens and show overlaps in a wide range of hosts and geographic distributions, they still exhibit skewed distributions in some ecological
and clinical niches. The larger clade includes strains that prefer tropics, carnivorous hosts, and occasionally lead to pandemic infections, whereas strains in the smaller clade show the opposite preferences as cold climate, herbivorous host, and often cause extra-intestinal and antibiotic-resistant infections. We further explore the reasons underlying these ecological distinctions by performing physiological experiments with our collection strains in the two clades. The strains in the larger clade show higher growth rate, especially in higher temperature, apparent vulnerability and poorer survival rates at heat stress due to faster growth, rapid response and motility to amino acid chemotaxis, as well as slower growth in polysaccharides carbohydrate sources when compared to the smaller clade (Fig.S3). We subsequently name the two clades as the Vig and the Slu clades, respectively.

**Figure 1.**
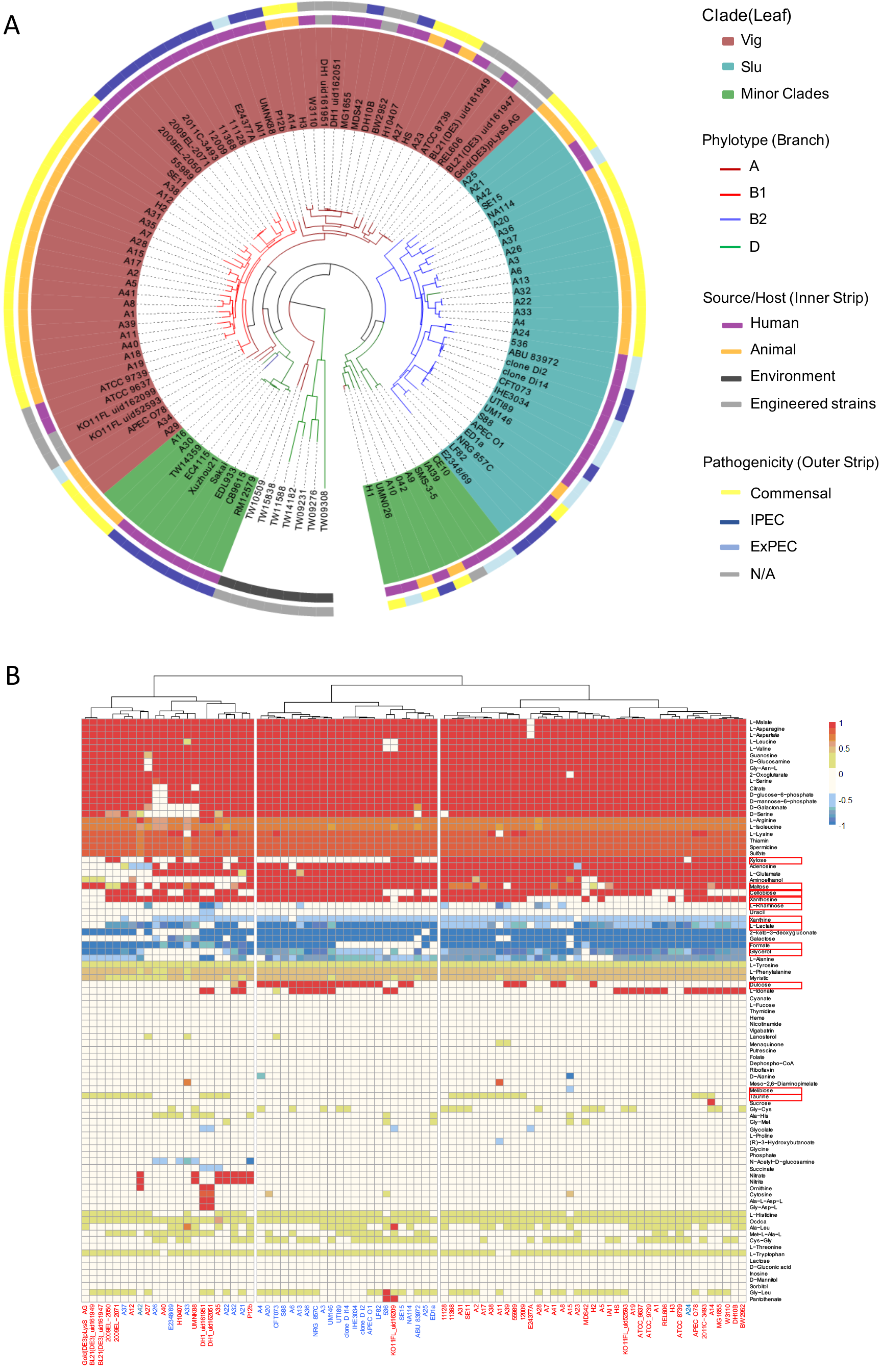
The two major *E*. *coli* clades: Vig and Slu. (A) An ML phylogeny based on the core-genome derived polymorphisms that separate host-related strains from the environmental strains (free-living in the tidal zone) and host-related strains into two major clades and a few minor clades. Names of each strain are labelled. Clade (colored leaf), phylotype (colored branch), origin (colored inner stripe), and pathotype (colored outer stripe) are indicated for each *E. coli* strain. (B) Prediction of metabolic compounds each strain uptakes and produces based on genome-scale models (GSMs) and genome annotations for the two clades. The cluster diagram represents 82 metabolic compounds with quantitative differences among strains, and those with significant inter-clade differences (p< 0.01, Wilcoxon test) are highlighted in red boxes. Each column represents a single strain with name labeled in red or blue color for the Vig or Slu, respectively. The top clustering is based on the Euclidean distance between strains and Ward clustering algorithms. Compound flux is normalized by its maximal value and colored in red for consuming and blue for producing (see the right scaling bar).

We examine genetic bases of the phenotypical distinctions between Vig and Slu in their genomes. First, we find apparent differences between the two clades in their profiles of dispensable genes based in a neighbor-joining tree (Fig. S4). By using two-sample Kolmogorov-Smirnov test to all dispensable genes, we identify 279 and 336 orthologs that are significantly enriched in Vig and Slu, respectively (Table S2). Half of these clade-associated-genes (CAGs) are metabolic genes according to the COG categories, and as expected (Fig.S5A), the Vig clade enriches more genes related to motility and metabolism of purine and amino acid than the Slu clade. Strains of the same clade share common patterns in dispensable metabolic genes regardless of host diet preferences (Fig.S5B). We further predict nutrient consumption and fermentation products by constructing the genome-scale models (GSMs) of metabolism for each strain *in silico* (17, 18), the two clades are largely separated by characteristic substrates and products, albeit a few confusions (Fig. 1B). Given a complete rich media which contains 16279 compounds, GSMs predict that our stains consume or produce 102 of these compounds, and 82 show differences between strains. A total of 12 compounds show significant difference (p<0.01, Wilcoxon test) between the two clades, i.e., the Slu strains uptake more plant-origin nutrients of dulcose, maltose, and cellobiose, whereas the Vig strains uptake more taurine and xylose. The Vig strains produce more formate, glycerol, and L-rhamnose, as compared to the Slu strains that produce more lactate and L-proline and are more active in xanthine metabolism. For dispensable virulence genes, the Vig pathogens are rich in T3SS and other secretion systems and the lethal Shiga-like toxins (Stx) (19), whereas the Slu pathogens have more genes for adhesion, invasion and Extended-Spectrum Beta Lactamases (ESBLs) that possibly facilitate their extra-intestinal infections and antibiotic-resistance (Fig.S5C,S5D). Next, we compare sequence variances between orthologs shared by the two clades. In the core genome, 126 and 227 non-synonymous polymorphisms in 97 and 168 genes show specificity for the Vig and Slu clades, respectively, which scatter along the core genome (Fig.S6A, Table S3). Half of the genes that contain these clade-associated variations (CAVs) are in the COG category of metabolism, which also underpin the metabolic distinctions between the two clades (Fig.S6B). Furthermore, homologues of the core genes involved in the metabolic pathways of amino acids and carbohydrates show a clear separation in their maximal-likelihood phylogeny when concatenated (Fig. S7). Dispensable orthologs with high prevalence (present in > 80% of the 104 strains) also show notable larger sequence distance in paired homologs between-clade than within-clade (Fig. S8). These differences in sequence distance suggest that each clade utilizes their own preferred alleles of both the core and dispensable genes, which are not readily traded between clades.

The obvious genetic differences between the clades suggest they exchange genetic material scarcely. We thus infer genome-wide recombination for each genome pair to see if recombination decreases sharply and shows signals of genetic separation between the two clades. Taking advantage of our complete genomes, we are able to infer recombination events by comparing synteny of aligned identical fragments. These fragments if moved to other locations are recognized as authentic recombinant fragments. Our result shows that extent of intra-clade recombination is rather extensive than that of inter-clade (Fig. 2A), in accordance with a previous report of less samples(20). For intra-clade genome pairs, the total lengths of recombinant fragments range 0.3Mb – 2Mb, and in some cases even approaching half of the genome, whereas those of inter-clade pairs are 10-fold shorter and never exceed 250kb (Fig. 2B). The intensity of intra-clade recombination exhibit almost the same extent even between commensal and pathogenic strains (Fig. 2B inlet), which implies that virulence genes can be freely transferred between commensals and pathogens of the same clade and makes efforts to distinguish them extremely difficult (21). Not only the intensity of recombination shows a clear cut-off between inter- and intra-clade recombination, but also logarithmically correlated to phylogenetic distance based on core genome polymorphism (Fig.2C) as previously estimated (22). Seemingly, the numbers of fragments that represent recombination frequency are much greater in intra-clade pairs than inter-clade pairs (Fig. 2D) and correlate with phylogenetic distance as well (Fig. 2E), except in case of very close strains where recombination frequency can be significantly underestimated due to the overlapping of recombinant fragments. The average size of recombinant fragments is also much longer in intra-clade genome pairs than what of inter-clade (Fig. 2F) and correlate with phylogenetic distance (Fig. 2G). Since restriction-modification system has been demonstrated to control recombination between species (23), we test its role in our samples with the DNA methylation information obtained from PacBio platform. However, the result shows no clade-specific restriction motif, although there are many strain-specific methylated motifs and a common motif of GATC in all *E. coli* strains (Fig. S9).

**Figure 2.**
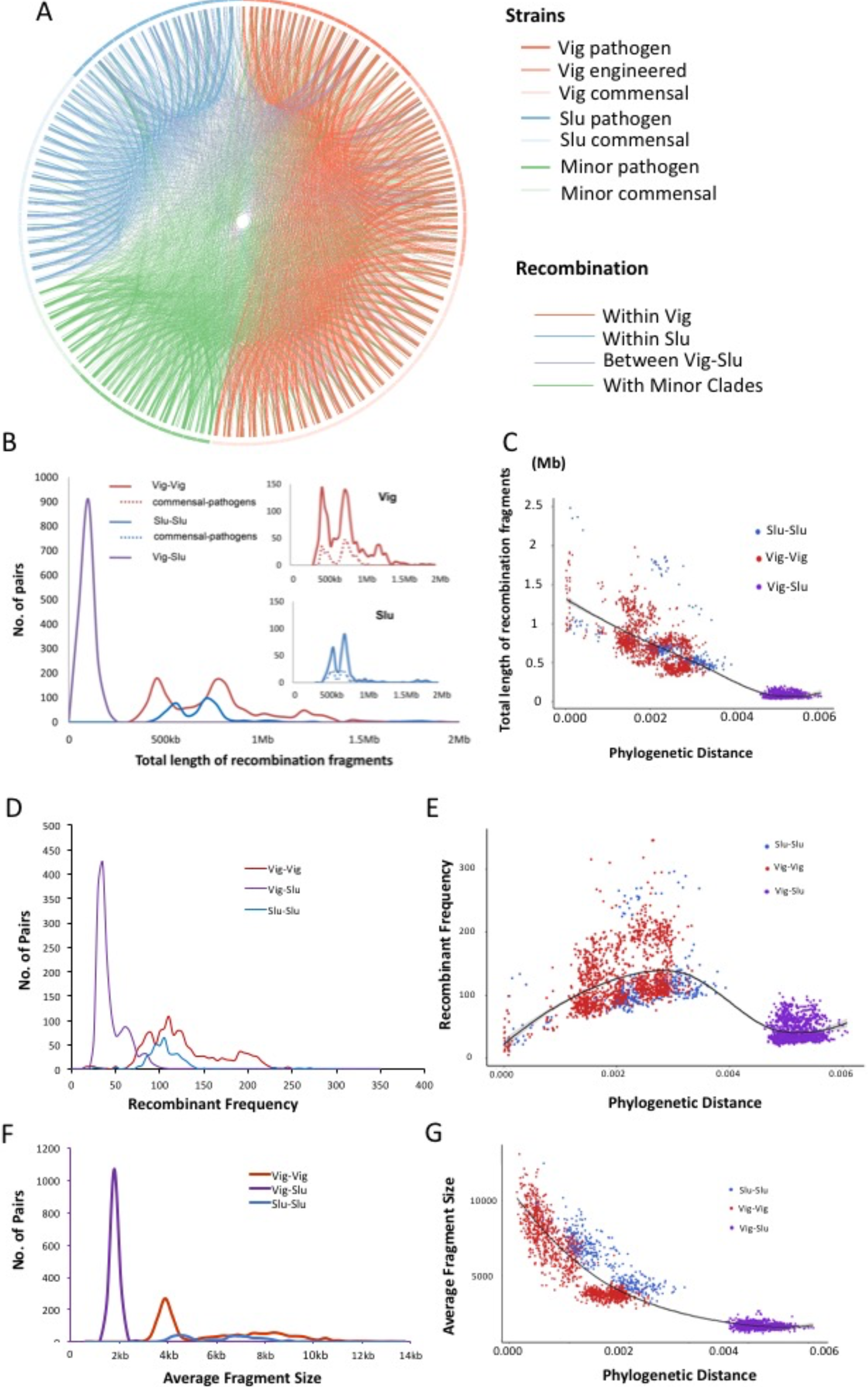
Intra- and inter-clade Recombinations. (A) Pair-wise assessment of recombination within and across clades. Strains are circularly displayed and the clades of Vig (red) and Slu (blue) are on both sides and the minor clade is placed (green) in the middle. Pathogenic and commensal strains are indicated in dark and light colors. Each line between strains indicates recombination between the two strains with thickness proportional to the total fragment length over 5kb. (B) Recombinant length distributions. Recombinants are grouped into intra-clade (Vig–Vig, red; Slu–Slu, blue) and inter-clade (Vig–Slu, purple), and those between commensal/pathogenic pairs (dotted line) are also shown in the insets. (C) Recombinant length is plotted as a function of phylogenetic distance. (D) A plot of recombination frequency (number of recombinant fragment). (E) Recombination frequency as a function of phylogenetic distance. (F) Average recombinant length distribution. (G) Average recombinant length as a function of phylogenetic distance.

The majority of recombinant fragments are in length of 3kb which fits in the common size of bacterial operons, even some super-fragments can approach 2Mb (Fig. 3A) as long as reported in other species (24, 25). As so efficient and frequent the recombination is in exchanging genetic materials especially within clade, we further examine its influence on the organization and metabolic scheme of the recipient genome. Firstly, we calculate the mobility of each single gene as its normalized frequency of being recombined. The result indicates a broad range of gene mobility from 0 to 100% and a clear negative correlation to prevalence, i.e., core genes which present in all strains are rather stable and seldom recombined, whereas dispensable genes, especially those only present in few strains, are actively transferred between strains (Fig. 3B). This is in accordance with the concept of Genome Organization Framework (GOF) (10), where core genes constitute a stable genome framework, leaving the intervals filled with mobile variable genes. Next, we examine the mobility of the clade-characteristic CAGs which show comparable overall mobility as other dispensable genes of similar prevalence (Fig. 3B), but differ their mobility in the two clades: Vig-CAGs are much more stable in Vig strains than in Slu strains, and those of Slu show opposite mobility in the two clades (Fig. 3C). As for the core genes containing CAVs, they are all rarely recombined, but still a little more stable in corresponding clades although the difference is not significant (Fig. 3D). The stability of core genes as well as CAGs in corresponding clades suggest that the recombination has little overall influences on genes with critical adaptive functions. Thus the extensive recombination does not disturb the framework of genome and adaptive metabolic scheme heavily, possibly due to strong negative selections for any changes that impair the fitness of host cell, according to the complexity hypothesis and cellular evolution(26).

**Figure 3.**
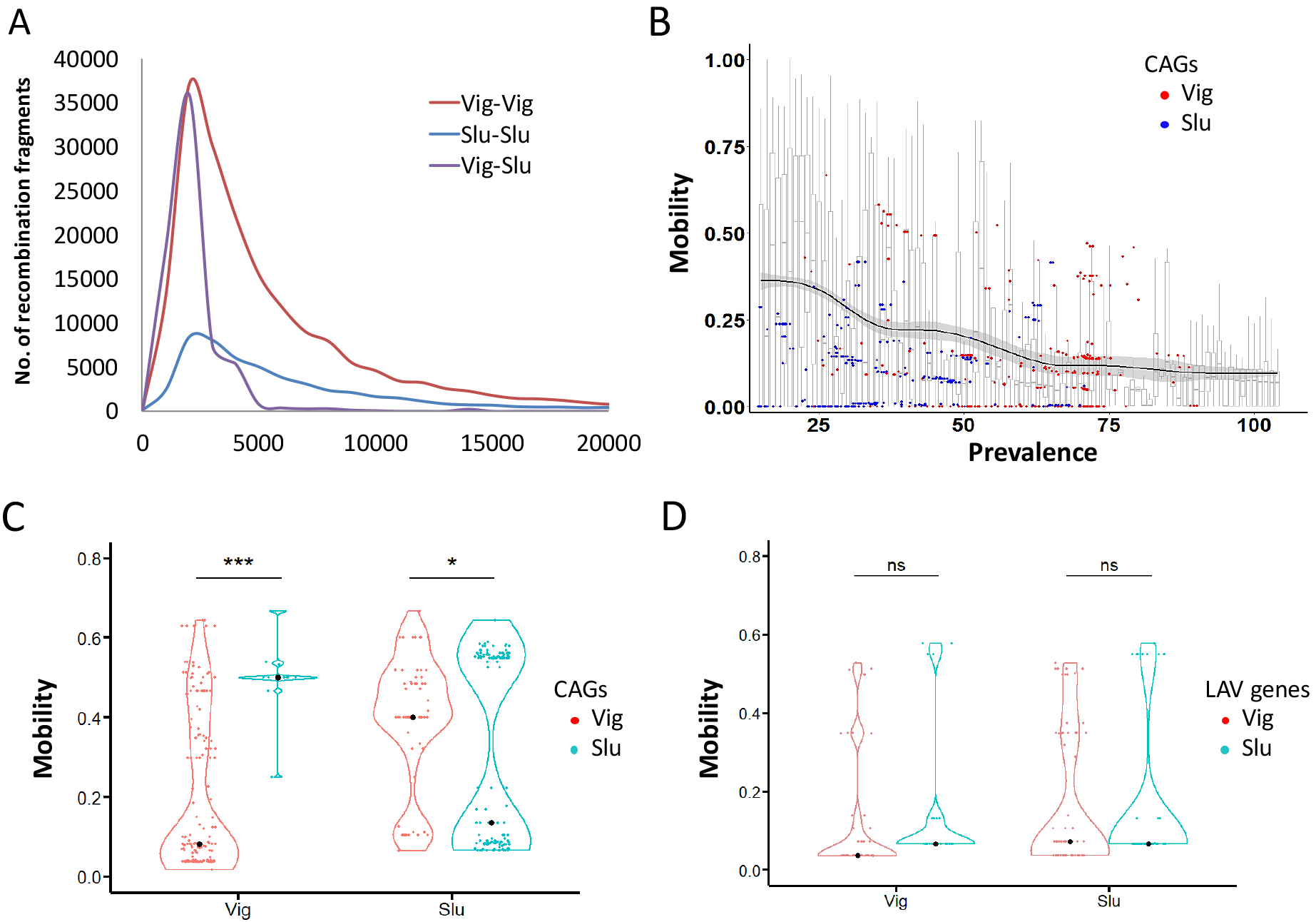
Gene mobility. (A) Recombinant size distributions. Recombinants of Vig intra-clade (Vig–Vig), Slu intra-clade (Slu– Slu), and inter-clade (Vig–Slu) are plotted as a function of size (bp) in red, blue, and purple, respectively. (B) Gene mobility (migration flexibility among genomes within and across clades) as a function of prevalence (the number of genomes that contain the gene). The regression curve (solid line) is based on generalized additive model with the shadow representing 95% confidence interval. CAGs (clade-associated genes) of Vig and Slu are indicated with red and blue solid circles, respectively. (C) Mobility of CAGs in Vig (red) and Slu (blue). (D) Mobility of CAV (clade-associated variations)-containing genes in Vig (red) and Slu (blue).

### Discussion

Our study is based on a dataset of high-quality complete genomes as well as physiological features experimentally acquired for the representative collection of *E. coli* strains. Two major clades of the population, Vig and Slu, show distinct ecological and clinical features due to their physiological differences and clade-featured metabolic profile. The phenotypes of the two clades are genetically underpinned by their differences in genome content and alleles they used for orthologous which make them adaptive to respective ecological niches. The thorough inference of recombination events for each pair of genomes indicates that recombination is rather frequent and intensive between within-clade strains but logarithmically reduced with phylogenetic distance, leading to genetic separation between clades. The extensive recombination, although keep cutting-and-pasting new genetic materials until they reach assortive genome locations, do not untidy GOFs and adaptive metabolic portfolios.

One of our findings is to quantitate the intensity of genetic recombination occurring within and across the *E. coli* clades we have defined, which is not easy due to weak signals of recombined sequences intrinsic to draft assemblies (11, 27). For a long time, recombination has been acknowledged to be very rare in asexual bacterial species, on which the theory of genome sweep has been established and genes adjacent to advantageous alleles may achieve high prevalence through genetic hitchhiking. Recent studies have questioned the theory and proposed bacteria as quasi-sexual populations where adaptive genes instead of genome sweep the population driven by nature selection (6, 7). Our data, providing exactly count on each recombination event for genome pairs, directly verify a high intra-clade recombination rate and quasi-sexual status of microbial populations.

The poor inter-clade recombination rate and its small recombinant fragment size suggest a mechanistic barrier in favor of speciation (28). In simulations of evolution model, given suitable mutation rate and enough population size, even without other involvement of nature selection or geographic separation, populations can diverge neutrally only if recombination rate depends on homolog distance (similar to phylogeny distance in our work) (29). In other words, if individuals in a population recombine more intensively with closer relatives, the disparity in recombination rate would by itself homogenize strains with similar genome sequences into a clade and build genetic barriers to others (28). As a result, the disparity of recombination intensity may consistently generate new clades; some of them are deemed to be adaptive or specialized to ecological niches and selected by nature and others may just drift on and off the edge. This is precisely the scenario of the Vig and Slu clade separation in our study, and such scenarios have been published in other bacteria, archaea (7, 30–33) and eukaryotes (34). However, molecular mechanism of the phylogeny-restricted recombination have yet to be described. Previous studies have proposed roles of geographic isolation, diversity of CRISPR-Cas system (35), DNA uptake signal sequences (36), and incompatible transfer mechanisms due to pili (37) for the mechanisms. Recently, high-efficiency recombination mechanisms, such as co-evolving phage and distributive conjugal transfer have been reported, which allows DNA fragments over 100kb in a single recombination event to happen but such a function deteriorates between remote relatives (20, 38). All these mechanisms with various transfer capacities and host range enable strains to utilize suitable means in exchanging genetic material according to phylogenetic distance of relatives.

The assessment of mobility for each single gene indicates that high recombination rate does not have impact on genes with critical adaptive functions as well as GOF and primary metabolic portfolios they built over long evolutionary time scale when genes or their alleles are constantly shuffled by the high-rate recombination until reach assortive locations and fixed by nature selection in a manner of cut-and-try. Besides niche adaptation, nature also selects for higher genetic compatibility among metabolic pathways, genes, and even alleles, such as the CAVs in our case, because conflicts among these components may undermine the competence of host strains in constant competitions with adjacent sister strains (39). Thus GOF and primary metabolic portfolios and their related genes may be resistant to recombination that results in allele substitution(40) and location rearrangement(41) because phylogenies with reduced integrated fitness may be eliminated by nature selections. Recent studies have shown genetic interaction and compatibility may create fitness cost and constrain the dissemination and evolution of dispensable genes (42,43), which similarly direct the transfer of genes and alleles intra- and inter-clade to confer Vig and Slu distinct ecological and clinical features. Thus recombination accelerates the optimization of the genome content to streamlines physiological functions and metabolic portfolios of host strains. High-rate recombination is able to help adaptive genes to sweep the population, which benefit recipient strains with competitive advantages and faster evolution.

In conclusion, our empirical data regarding to the separation of clades Vig and Slu in species *E. coli* clarify the role of recombination in population diversification and evolution. The extensive intra-clade recombination interrupts linkage between genes and leaves them independent under selections, but do not disarranged the optimized GOF and metabolic scheme of host strains possibly due to critical selection for streamlined physiological activities. Meanwhile, the extensive recombination within a clade coheres strains into a quasi-sexual population, but its intensity declines sharply as phylogeny distance increases, which builds barriers between clades and diverge the population.

### Methods and Materials

#### Strains and MLST typing

A world-wide collection of 202 *E. coli* strains from vertebrate hosts (13) was kindly provided by Professor Shulin Liu (Genomics Research Center, Harbin Medical University, Harbin, China). A total of 149 successfully resuscitated strains were subjected to MLST sequencing and construction of a maximum-likelihood phylogeny using RaxML with model GTR and 1,000 bootstraps, together with 59 complete genomes of human-host deposited in NCBI (ftp://ftp.ncbi.nlm.nih.gov/genomes/Bacteria/) as of July 2013 and seven draft genomes published by a study of environmental strains (14) from ftp://ftp.ncbi.nlm.nih.gov/genomes/Bacteria_DRAFT/.

#### Genome sequencing, annotation, and phylogeny inference

The 45 selected representative strains were sequenced by using Hiseq 2000 and a paired-end sequencing protocol (2 × 100 bp run) at depth of 150×. Long reads are also added for contiguity from PacBio RS II with one or two cells per strain. After quality-filtration using the SMRT Analysis software suite (Pacific Biosciences Inc.), *de novo* assembly was performed according to the Hierarchical Genome Assembly Process (HGAP) workflow, and consensus sequences were corrected using Pilon (v1.22) with the clean reads generated from Illumina Hiseq 2000. Genomes were annotated with GeneMark.hmm and BLAST against the COG database and KEGG GENES (www.genome.jp/kegg/genes.html). Orthologs were identified by using PGAP (44), and the annotated genomes show apparent consistency when compared to the genome framework generated from GAAP (http://gaap.big.ac.cn), a synteny of core genes on genomic scale (45). Phylotypes of all strains were identified *in silicon* according to the presence of three phylotype-specific genes or fragments as previously described (46).

#### Measurement of growth rate, survival rate, and mobility

Refer to SI.

#### Genome content similarity, gene distribution, and homologue distance between the Vig and Slu clades

The Bray–Curtis dissimilarity index d was calculated for paired genomes as 1 − [2 * Sij/(Si + Sj)], where Sij is the number of genes shared by strains i and j, and Si, Sj as that in strains i and j, respectively. Two-sample Kolmogorov-Smirnov test was applied to select ortholog significantly enriched in Vig or Slu, when *p* <0.01. Antibiotic-resistance and Shiga-like toxin genes were identified by BLAST against CBMAR database (http://14.139.227.92/mkumar/lactamasedb) and the protein sequences of the toxin from *E. coli* O157:H7 (lcl|AB015056.1_prot_BAA88123.1_1 for A-subunit and lcl|AB015056.1_prot_BAA88124.1_2 for B-subunit). Pairwise distance of orthologs was calculated by using Protdist in the Phylip package.

#### Metabolism reconstruction

The metabolic profile of each strain was reconstructed *in silico* using the ge-nome-scale models (GSMs) as described previously (17). Briefly, we first removed the pseudogenes in annotation and used ModelSEED (http://modelseed.org/) to build the genome-scale models based on flux balance analysis (18). Here we used the condition of “gram negative template” and “complete media” which contains 16279 compounds in KBase (ftp://ftp.kbase.us/assets/KBase_Reference_Data/Biochemistry). A compound was predicted to be consumed or produced when corresponding transport, metabolism or biosynthesis reactions were available in a strain.

#### Recombination inference

To infer genome-wide recombination, we utilized gmos (Genome MOsaic Structure) to make local alignments between paired genomes and identify identical fragments over 600bp (47) as candidate recombinant fragments. We further used the syntenic information of these candidates which parsimoniously retain those changed locations as authentic recombination fragments. Gene mobility was calculated as the percentage of being recombined in total genome pairs the gene was present. The recombination of a gene was also identified using the syntenic information as what for recombinant fragments.

## Data Availability

The genome sequences we contribute are available on the NCBI GenBank (accession No. CP012760, CP012781, CP029328, CP029327, CP028732-CP028772) and Genome Sequence Archive of GSA database (http://gsa.big.ac.cn, Accession No. GWHAACT-GWHAAEJ). Raw data of sequencing are deposited in the GSA database (PacBio, Accession No. CRR028990-CRR029033; Hiseq, Accession No. CRR019978-CRR020022).

## Competing interests

The authors declare no competing interests.

## Funding

This work was supported by the National Natural Scientific Foundation of China [31470180, 31471237, 31671350and 30971610], the National Key Research and Development Program of China [2016YFC0903800], Programs of Ministry of Health of the People’s of China [201402018], Programs of the Chinese Academy of Sciences [QYZDY-SSW-SMC017, YZ201568, YZ201402], Programs of Beijing Municipal Science and Technology Project [Z171100001317011], Taicang Municipal Science and Technology Project[TC2016SF08].

## Author contribution

J.Y. and Y.K. conceived the project and led the writing; L.Y., Y.C., R.F., and J. W. compiled the data; and X.S., F.C., Z.H., Z.G., X.J., Q.L, Q.M., J.W., J.X., and S.H. analyzed the data. All authors contributed to the writing and/or intellectual development of the manuscript.

## References

1. Fraser C, Hanage WP, & Spratt BG (2007) Recombination and the nature of bacterial speciation. Science (New York, N.Y.) 315(5811):476-480.

2. Shapiro BJ, Leducq JB, & Mallet J (2016) What Is Speciation? PLoS genetics 12(3):e1005860.

3. Cohan FM (2001) Bacterial species and speciation. Systematic biology 50(4):513-524.

4. Cohan FM & Perry EB (2007) A systematics for discovering the fundamental units of bacterial diversity. Current biology: CB 17(10):R373-386.

5. Bendall ML, et al. (2016) Genome-wide selective sweeps and gene-specific sweeps in natural bacterial populations. The ISME journal 10(7):1589-1601.

6. Rosen MJ, Davison M, Bhaya D, & Fisher DS (2015) Microbial diversity. Fine-scale diversity and extensive recombination in a quasisexual bacterial population occupying a broad niche. Science (New York, N.Y.) 348(6238):1019-1023.

7. Shapiro BJ, et al. (2012) Population genomics of early events in the ecological differentiation of bacteria. Science (New York, N.Y.) 336(6077):48-51.

8. Doolittle WF (2012) Population genomics: how bacterial species form and why they don’t exist. Current biology: CB 22(11):R451-453.

9. Fraser C, Alm EJ, Polz MF, Spratt BG, & Hanage WP (2009) The bacterial species challenge: making sense of genetic and ecological diversity. Science (New York, N.Y.) 323(5915):741-746.

10. Kang Y, et al. (2014) Flexibility and symmetry of prokaryotic genome rearrangement reveal lineage-associated core-gene-defined genome organizational frameworks. MBio 5(6):e01867.

11. Ravenhall M, Skunca N, Lassalle F, & Dessimoz C (2015) Inferring horizontal gene transfer. PLoS computational biology 11(5):e1004095.

12. Martinez-Castillo A & Muniesa M (2014) Implications of free Shiga toxin-converting bacteriophages occurring outside bacteria for the evolution and the detection of Shiga toxin-producing Escherichia coli. Frontiers in cellular and infection microbiology 4:46.

13. Souza V, Rocha M, Valera A, & Eguiarte LE (1999) Genetic structure of natural populations of Escherichia coli in wild hosts on different continents. Applied and environmental microbiology 65(8):3373-3385.

14. Luo C, et al. (2011) Genome sequencing of environmental Escherichia coli expands understanding of the ecology and speciation of the model bacterial species. Proceedings of the National Academy of Sciences of the United States of America 108(17):7200-7205.

15. Didelot X, Meric G, Falush D, & Darling AE (2012) Impact of homologous and non-homologous recombination in the genomic evolution of Escherichia coli. BMC genomics 13:256.

16. Kaas RS, Friis C, Ussery DW, & Aarestrup FM (2012) Estimating variation within the genes and inferring the phylogeny of 186 sequenced diverse Escherichia coli genomes. BMC genomics 13:577.

17. Monk JM, et al. (2013) Genome-scale metabolic reconstructions of multiple Escherichiacoli strains highlight strain-specific adaptations to nutritional environments. Proceedings of the National Academy of Sciences of the United States of America 110(50):20338-20343.

18. Henry CS, et al. (2010) High-throughput generation, optimization and analysis of genome-scale metabolic models. Nature biotechnology 28(9):977-982.

19. Tarr PI, Gordon CA, & Chandler WL (2005) Shiga-toxin-producing Escherichia coli and haemolytic uraemic syndrome. Lancet 365(9464):1073-1086.

20. Dixit PD, Pang TY, Studier FW, & Maslov S (2015) Recombinant transfer in the basic genome of Escherichia coli. Proceedings of the National Academy of Sciences of the United States of America 112(29):9070-9075.

21. Picard B, et al. (1999) The link between phylogeny and virulence in Escherichia coliextraintestinal infection. infection and immunity 67(2):546-553.

22. Roberts MS & Cohan FM (1993) The effect of DNA sequence divergence on sexual isolation in Bacillus. Genetics 134(2):401-408.

23. Pleska M, et al. (2016) Bacterial Autoimmunity Due to a Restriction-Modification System. Current biology: CB 26(3):404-409.

24. Boritsch EC, et al. (2016) Key experimental evidence of chromosomal DNA transfer among selected tuberculosis-causing mycobacteria. Proceedings of the National Academy of Sciences of the United States of America 113(35):9876-9881.

25. Chen L, Mathema B, Pitout JD, DeLeo FR, & Kreiswirth BN (2014) Epidemic Klebsiellapneumoniae ST258 is a hybrid strain. mBio 5(3):e01355-01314.

26. Woese CR (2002) On the evolution of cells. Proceedings of the National Academy of Sciences of the United States of America 99(13):8742-8747.

27. Nielsen KM, Bohn T, & Townsend JP (2014) Detecting rare gene transfer events in bacterial populations. Frontiers in microbiology 4:415.

28. Dixit PD, Pang TY, & Maslov S (2017) Recombination-Driven Genome Evolution and Stability of Bacterial Species. Genetics 207(1):281-295.

29. Falush D, et al. (2006) Mismatch induced speciation in Salmonella: model and data.Philosophical transactions of the Royal Society of London. Series B, Biological sciences 361(1475):2045-2053.

30. Didelot X, et al. (2011) Recombination and population structure in Salmonella enterica. PLoS genetics 7(7):e1002191.

31. Huang CL, et al. (2015) Ecological genomics in Xanthomonas: the nature of genetic adaptation with homologous recombination and host shifts. BMC genomics 16:188.

32. Zwick ME, et al. (2012) Genomic characterization of the Bacillus cereus sensu lato species: backdrop to the evolution of Bacillus anthracis. Genome research 22(8):1512-1524.

33. Cadillo-Quiroz H, et al. (2012) Patterns of gene flow define species of thermophilic Archaea. PLoS biology 10(2):e1001265.

34. Ellison CE, et al. (2011) Population genomics and local adaptation in wild isolates of amodel microbial eukaryote. Proceedings of the National Academy of Sciences of the United States of America 108(7):2831-2836.

35. Samson JE, Magadan AH, & Moineau S (2015) The CRISPR-Cas Immune System and Genetic Transfers: Reaching an Equilibrium. Microbiology spectrum 3(1):Plas-0034-2014.

36. Frye SA, Nilsen M, Tonjum T, & Ambur OH (2013) Dialects of the DNA uptake sequence in Neisseriaceae. PLoS genetics 9(4):e1003458.

37. Cehovin A, et al. (2013) Specific DNA recognition mediated by a type IV pilin. Proceedings of the National Academy of Sciences of the United States of America 110(8):3065-3070.

38. Gray TA, Krywy JA, Harold J, Palumbo MJ, & Derbyshire KM (2013) Distributive conjugal transfer in mycobacteria generates progeny with meiotic-like genome-wide mosaicism,allowing mapping of a mating identity locus. PLoS biology 11(7):e1001602.

39. Bao YJ, Shapiro BJ, Lee SW, Ploplis VA, & Castellino FJ (2016) Phenotypic differentiation of Streptococcus pyogenes populations is induced by recombination-driven gene-specific sweeps. Scientific reports 6:36644.

40. Kacar B, Garmendia E, Tuncbag N, Andersson DI, & Hughes D (2017) Functional Constraints on Replacing an Essential Gene with Its Ancient and Modern Homologs. MBio 8(4).

41. Ma Q, et al. (2013) Computational analyses of transcriptomic data reveal the dynamic organization of the Escherichia coli chromosome under different conditions. Nucleic acids research 41(11):5594-5603.

42. San Millan A, Toll-Riera M, Qi Q, & MacLean RC (2015) Interactions between horizontally acquired genes create a fitness cost in Pseudomonas aeruginosa. Nature communications 6:6845.

43. Porse A, Schou TS, Munck C, Ellabaan MMH, & Sommer MOA (2018) Biochemical mechanisms determine the functional compatibility of heterologous genes. Nature communications 9(1):522.

44. Zhao Y, et al. (2012) PGAP: pan-genomes analysis pipeline. Bioinformatics 28(3):416-418.

45. Yuan L, et al. (2017) GAAP: Genome-organization-framework-Assisted Assembly Pipeline for prokaryotic genomes. BMC genomics 18(Suppl 1):952.

46. Clermont O, Bonacorsi S, & Bingen E (2000) Rapid and simple determination of the Escherichia coli phylogenetic group. Applied and environmental microbiology 66(10):4555-4558.

47. Domazet-Loso M & Domazet-Loso T (2016) gmos: Rapid Detection of Genome Mosaicism over Short Evolutionary Distances. PloS one 11(11):e0166602.

